# Proteomic comparison of Marburg and Kasokero virus infection in natural host Egyptian rousette bat reveals distinct antiviral pathways

**DOI:** 10.64898/2026.09.28.755161

**Authors:** Brooke N. Genovese, Nistara Randhawa, Benjamin A. Neely, Gabriela Grigorean, Amy J. Schuh, Brian R. Amman, Jessica A. Elbert, Simon J. Anthony, Jonna A.K. Mazet, Jonathan S. Towner, Brian H. Bird

## Abstract

Bats are natural reservoir hosts for numerous zoonotic viruses, yet the molecular mechanisms enabling viral persistence without overt disease remain incompletely understood. The Egyptian rousette bat (Rousettus aegyptiacus, ERB) is the sole known natural reservoir of Marburg virus (MARV) and a key host in the enzootic cycle of the tick-borne Kasokero virus (KASV), providing a unique system to compare host responses to different RNA viruses within the same species. Here, we applied serial cross-sectional serum proteomics integrated with tissue-specific viral kinetics to characterize systemic host responses to experimental MARV and KASV infection in captive-reared bats. Nearly 16 % of detected proteins (67/419) were found exclusively in infected animals, yet canonical acute inflammatory signatures driving pathology in other susceptible hosts were absent (e.g., in humans and non-human primates). Differential abundance and detection analyses identified both shared and virus-dependent responses, with MARV infection eliciting limited and transient perturbations, while KASV induced broader and more sustained engagement of complement, lectin pathway, and hepatometabolic proteins. Network-based analysis uncovered coordinated proteasome modules — including circulating immunoproteasome complexes — consistent with enhanced antigen-processing capacity is a feature of these virus-host dynamics. Together, these findings reveal how ERBs mount structured, virus-dependent systemic responses, offering new insight into mechanisms underlying viral infection in natural reservoir hosts.

## Introduction

Bats (Order: Chiroptera) comprise one of the most diverse mammalian taxa, providing core ecosystem services such as pollination, seed dispersal, generation of nutrient-rich, organic manure (e.g., bat guano), and arthropod suppression(1, 2). As with many highly diverse vertebrate groups, bats harbor a broad range of zoonotic viruses, including paramyxoviruses, coronaviruses, and filoviruses, typically without overt signs of illness(3, 4). Among bat–virus systems, infections of the Egyptian rousette bat (Rousettus aegyptiacus; ERB) are one of the most intensively studied. ERBs are the only known natural reservoir of Marburg virus (MARV; family Filoviridae), the causative agent of severe and often fatal Marburg virus disease in humans and non-human primates(5, 6). ERBs are also the sole known wildlife host in the enzootic cycle of Kasokero virus (KASV; family Nairoviridae), a zoonotic tick-borne virus maintained primarily through interactions with the argasid tick Ornithodoros (Reticulinaeus) faini within bat colonies(7–9). These dual associations position ERBs as a uniquely informative system for examining how distinct RNA viruses interact with their reservoir host.

Although MARV and KASV both produce clinically inapparent infections in ERBs, they differ markedly in their within-host dynamics and transmission ecology. MARV infection produces a brief viremia (mean duration ≈ 3 days post-infection [DPI]), whereas oral shedding may persist for up to three weeks(10–12). Bat-to-bat transmission, potentially mediated by biting, is thought to sustain viral circulation within roosts(10). In contrast, KASV infection is characterized by an early burst of hepatic viral replication, self-limiting lymphohistiocytic hepatitis(13), longer duration of viremia (mean ≈ 7 DPI), and detectable oral shedding from 2 to 14 DPI (mean duration ≈ 8 DPI(8)). Recent experimental work further shows that KASV coinfection can alter MARV infection kinetics in ERBs, increasing viral loads, prolonging shedding, and elevating the probability of supershedding individuals(14).

Transcriptomic studies of MARV-infected ERBs show induction of interferon-stimulated genes (ISGs) alongside limited expression of classical pro-inflammatory mediators(15). Complementary genomic analyses of ERB cells and tissues have identified the absence of proteins associated with prototypic acute-phase responses (pentraxins e.g., C-reactive protein, serum amyloid P)(16) and expansions in Natural Killer cell receptors, MHC class I genes, and type I interferons(17). Together, gene-expression studies suggest that ERBs control filovirus infection through tightly regulated antiviral responses without strong inflammatory pathology(18–20).

Importantly, filovirus-mediated disease severity is driven by the interaction between host responses and viral replication dynamics, rather than from intrinsic properties of the virus alone(21). This is illustrated by experiments in ERBs, where suppressing early inflammatory signaling with a potent steroid (dexamethasone) resulted in uncontrolled MARV replication and severe hepatic pathology resembling disease observed in humans(20). In this framework, apparent disease tolerance is best understood as an emergent property of coordinated host–virus interactions rather than a simple absence of immune activation.

Major gaps remain in our collective understanding of ERB immunobiology. Classical immunological methods are constrained by the scarcity of species-specific reagents(22), and transcriptomic analyses provide only indirect insight into functional immune activity. Critically, many antiviral processes — including interferon signaling, cytotoxic lymphocyte function, and innate immune pathways targeted by viral immune-evasion mechanisms — are regulated extensively at post-transcriptional and post-translational levels (23–25). As a result, transcriptional changes may not necessarily reflect protein abundance or functional activity. Proteomic approaches therefore provide a needed complement to transcriptomic studies, enabling direct measurement of host proteins and the molecular pathways that govern infection outcomes.

Building on our initial work to characterize the serum proteome of clinically-healthy, non-experimentally infected, captive-reared ERBs(26), we combined serial cross-sectional serum proteomics with multi-tissue viral kinetics to characterize host responses to MARV and KASV infection in ERBs. Using global proteomic profiling, differential abundance analyses, and network-based co-abundance modeling, we identify both shared and virus-dependent proteomic signatures across tissues and throughout infection. A key strength of this study is the use of low-passage, bat- and tick-derived viral isolates (371bat MARV; UGA-Tick-20170128) integrated with previously published tissue-level viral replication data from the same animals(8, 10). These analyses reveal coordinated protein networks associated with antiviral defense, immune regulation, and cellular homeostasis, providing new insight into how different viruses engage the same reservoir host.

## Results

We analyzed the serum proteome of a total of 30 captive-reared ERBs from two previous experimental infection and serial euthanasia studies describing viral kinetics for MARV (n=15)(10) and KASV (n=15)(8) (Fig. 1). Infection profiles were compared to uninfected bats (n=6) and matched days post-infection (DPI) between MARV and KASV groups (see Methods). In both infection studies, animals were humanely euthanized, and necropsies were performed at defined timepoints post-infection to assess infection dissemination across different tissues.

**Figure 1.**
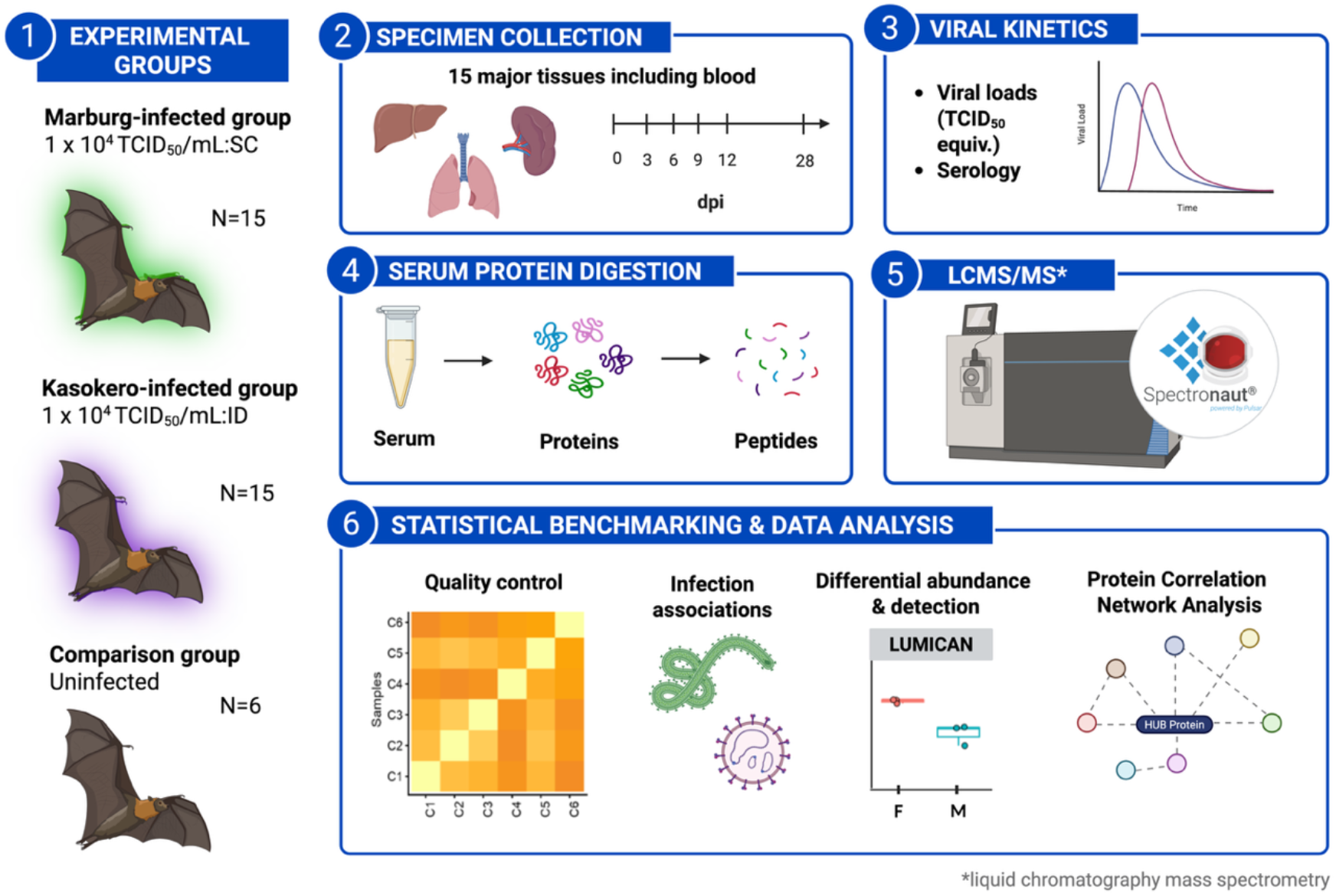
Experimental overview. Blood specimens were obtained from the cephalic vein of adult, captive-reared, non-pregnant Egyptian rousette bats experimentally-infected with either MARV (n=15) or KASV (n=15). Specimens from an additional six uninfected bats were included for comparison. The serum samples were gamma-irradiated prior to performing protein digestion by the S-trap method (see ‘Methods’ for details). From here, the samples (peptides) were analyzed by “shotgun” liquid chromatography mass-spectrometry (LC-MS/MS) operated in data-independent acquisition mode (DIA) and subsequently processed using Spectronaut 18. The resulting Spectronaut report, which included quantitative peptide and protein data, was the primary input for downstream bioinformatic analyses in R.

### Virus infection induces changes in the peripheral proteome

After filtering for low-confidence peptide matches, we quantified 4,814 peptides corresponding to 419 proteins across all 36 individuals. Unsupervised global analyses demonstrated clear clustering of uninfected samples, as well as clustering by infection type (Fig. 2A-B) and days post-infection (DPI) (Supplemental Fig. 1). Variance partitioning revealed that group (infection status x DPI) was the primary driver of protein abundance variation, explaining 23.1 % of variance on average (median: 20.1%), while sex and batch effects each contributed minimally (1 to 2 % of variance) (Supplementary Fig. 2).

**Figure 2.**
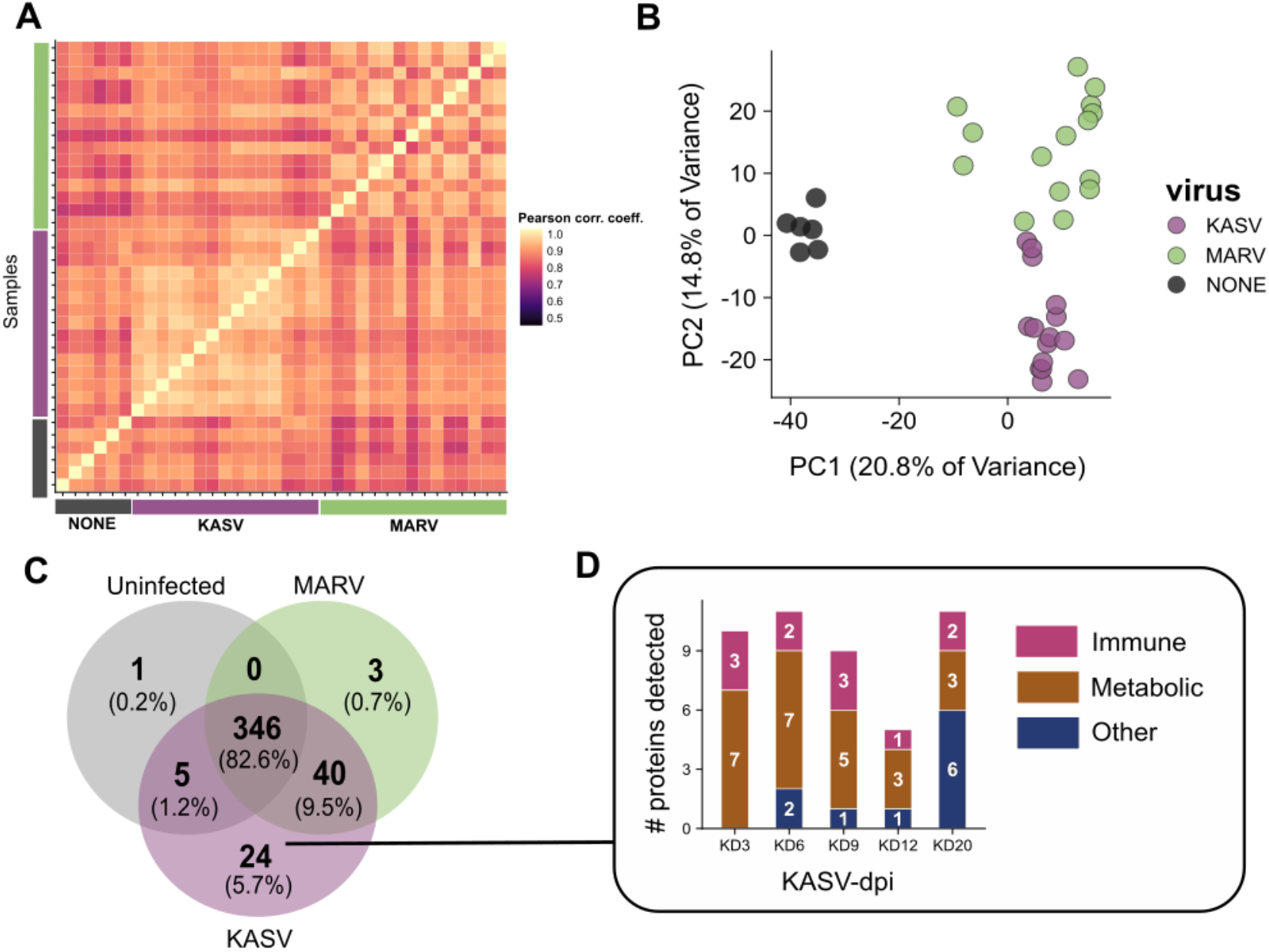
Global overview of serum protein profiles in uninfected and virus-infected bats. **A) Sample correlation heatmap**. A correlation heatmap among samples with Pearson correlation coefficients (r) on a scale of 0.5 to 1.0. A correlation coefficient of 1 indicates a perfect positive linear correlation between sample pairs. Samples from uninfected bats are in grey and labeled ‘NONE’, MARV-infected bats are shown in green, and KASV-infected bats are in purple. This color scheme is applied throughout the figure panel. All samples are grouped by sequential time point within their infection groups. **B) Principal Component Analysis (PCA) plots by infection (virus) type.** The plot shows protein profiles of sample points colored by infection type and variance of PC1 (x-axis) and PC2 (y-axis). **C) Commonly and exclusively quantitated proteins across uninfected and infected (MARV or KASV) groups.** Counts are out of a total 419 individual quantitated proteins across all samples. The only protein found solely in uninfected samples was alpha-hemoglobin stabilizing protein (AHSP), and it was consistently detected in all six uninfected bats. **D) Temporal detection counts of 24 KASV-exclusive proteins.** The y-axis represents number of proteins as counts, and the x-axis represents each day post-infection (dpi) time point (e.g. KD = KASV dpi). Broad functional categories are shown in pink (immune), brown (metabolic), and other (dark blue).

During MARV and KASV infection, the circulating proteome of infected bats shared 80 % (346/419) of detected proteins with uninfected bats (Fig. 2C). Comparative analysis of the 40 proteins detected in both MARV and KASV infections revealed virus-dependent differences in detection frequency across multiple functional categories (Supplementary Fig. S4), with differentially detected proteins distributed broadly rather than enriched within a single pathway. Exploratory enrichment analysis of these 40 proteins found in both MARV and KASV infection groups identified “Antigen Processing and Presentation of Peptide Antigen via MHC Class I (GO:0002474)” within the top ontology biological process (GO:BP) terms, linked to the proteins B2M, HLA-A, HSP90B1, HSPE1, and PDIA3.

More than 35 % (24/67) of all infection-exclusive proteins were detected only in KASV-infected bats across all five time points and were predominantly metabolic enzymes (n=13), particularly during acute infection (3 DPI to 6 DPI) (Fig. 2C-D; Supplementary Fig. S3). Immune-related proteins uniquely detected in KASV infection included ARGI1, ASS1, CSF1R, CD9 (20 DPI only), and a distinct CAMP-like protein (A0A7J8HSR3; CAMP2) detected at 6, 12, and 20 DPI. Low sequence identity ( ≈28 %) and extensive non-overlapping regions between CAMP2 and the second CAMP-like protein support their classification as distinct CAMP family members rather than splice variants (Supplementary Fig. S5). Full protein lists are provided in the Supplementary Materials.

### Near-complete constitutive proteasome detected in virus-infected bat serum

Given the enrichment of antigen presentation–related proteins in initial detection analyses, we examined proteasome subunit representation prior to global network modeling. Sixteen of the 17 canonical subunits were detected, consistent with a near-complete constitutive 20S proteasome in serum from KASV- and MARV-infected ERBs (Fig. 3A-B). Detected components included nearly all α-subunits, catalytically active and inactive β-subunits, the inducible immunoproteasome subunits PSMB8 and PSMB10, and the 19S regulatory subunit PSMD2, consistent with circulating 26S proteasome components. Peptide-level analysis confirmed mature PSMB10 detection across samples, whereas the constitutive subunit PSMB7 was consistently detected across infection time points and most uninfected bats (5/6).

**Figure 3.**
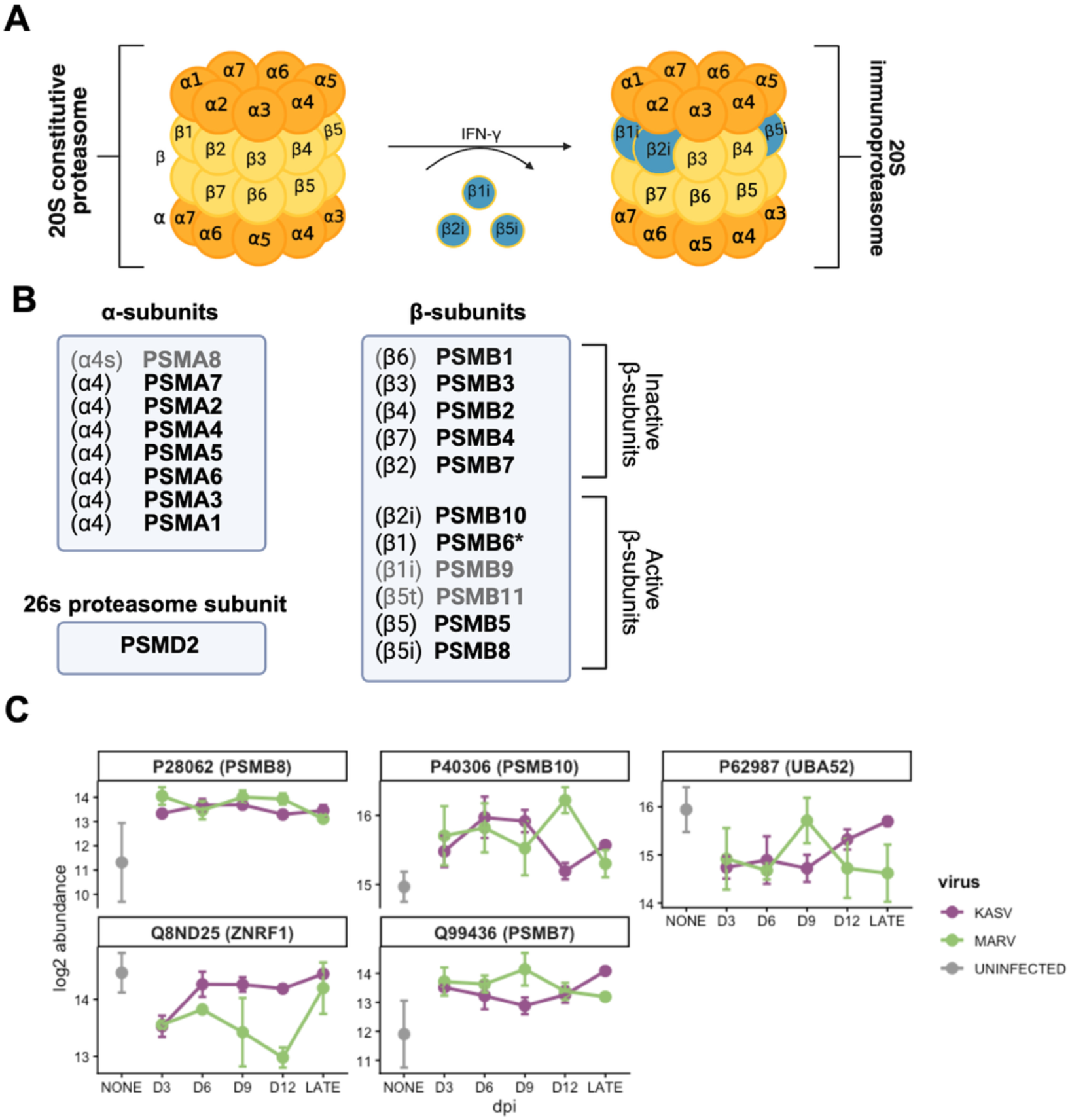
Serum proteasome profiles in infected ERBs. A) Architecture of the eukaryotic 20S core particle proteasome and immunoproteasome. The α-rings are shown in orange and the inner β-rings are shown in yellow. Subunit substitution delineates immunoproteasome specificity: interferon (IFN)-γ exposure induces substitution of constitutive β1/β2/β5 subunits with inducible β1i (PSMB9; LMP2), β2i (PSMB10; MECL-1), and β5i (PSMB8; LMP7). **B) Lists of detected proteasome subunits**. The names of the proteasome subunits are mentioned following the common nomenclature (α and β). Subunits that were detected in this dataset are in black, while subunits that were not detected in this dataset (including in uninfected bats) are shown in grey. **C) Kinetic profiles of proteasomal subunits and ubiquitin-related proteins in ERB serum throughout virus infection.** Line plots represent log2 abundance values of five key proteins identified in serum specimens from uninfected (grey; labeled ‘NONE’), KASV-infected (purple), and MARV-infected (green) ERBs. Data points represent the mean abundance at each day post-infection (dpi), with error bars indicating the standard error of the mean. Bat proteins are listed by the UniProt ID of the corresponding human ortholog alongside the human gene symbol. Note: * = detected only in MARV-infected ERBs

Building on the detection of near-complete circulating 20S proteasome and immunoproteasome subunits, we identified additional evidence of coordinated ubiquitin– proteasome system (UPS) activation in both infected cohorts (Fig. 3C). Immunoproteasome subunits PSMB8 and PSMB10 were elevated at 3 DPI relative to uninfected bats, whereas the constitutive subunit PSMB7 remained stable, with all three trending toward baseline at later time points. In contrast, the E3 ligase ZNRF1 and ubiquitin precursor UBA52 were consistently reduced following both infections. Circulating HLA-A, which functions in MHC class I antigen presentation following proteasomal peptide processing, was detected sporadically in infected bats but not in uninfected bats (Supplementary Table 2).

### Differential abundance analysis of serum proteins reveals time- and virus-dependent changes in the serum proteome

To further investigate the proteomic alterations that characterize each stage of infection, we next focused on the proteins that were significantly increased or decreased in abundance. Differential abundance analysis of 288 consistently quantified serum proteins revealed time- and virus-dependent remodeling of the circulating proteome (Fig. 4A). Comparisons between uninfected and MARV-infected bats showed limited perturbation at most time points, with few proteins reaching significance (q ≤ 0.05) and modest effect sizes. The strongest response occurred at 6 DPI, where 33 proteins were differentially abundant (13 up-regulated; 20 down-regulated). Preliminary ontological analysis at this peak time point indicated enrichment of immune processes, including humoral immune responses, among upregulated proteins, whereas downregulated proteins were associated with anatomical structure development and organelle biogenesis (Fig. 4B).

**Figure 4.**
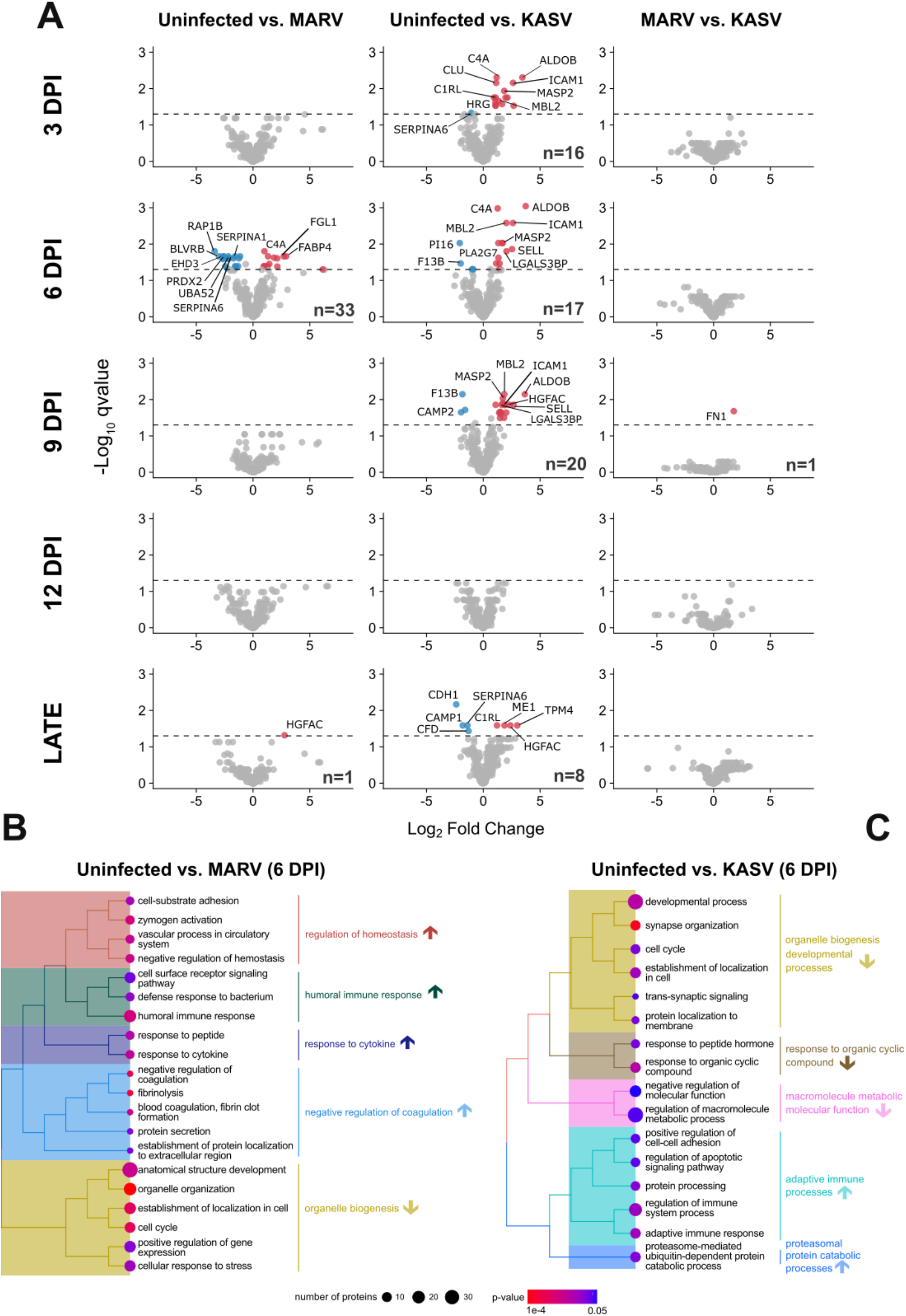
A) Differential abundance of serum proteins following MARV and KASV infection. Volcano plots depict DAA results across 288 proteins common to uninfected and infected (MARV and KASV) groups. For each contrast "A vs B," fold changes represent B/A. For example, in "Uninfected vs MARV," a positive log₂ fold change indicates higher protein abundance in MARV samples. Positive log₂ fold changes are shown in red; negative log₂ fold changes in blue. Adjusted P-values (i.e., q-values) derived from pairwise contrasts of the linear regression model are adjusted for sex and batch. The total number of statistically significant proteins (q < 0.05) per comparison is shown in the bottom right corner of each plot, and the top ten proteins with the smallest q-values are labeled by their human gene symbol. **B-C) GSEA biological trend analysis of host response in MARV and KASV infections**. Treeplots illustrate exploratory biological trends in serum protein responses in (B) Uninfected vs. MARV (6 DPI) and (C) Uninfected vs. KASV (6 DPI) comparisons. Due to the focused nature of the protein panel, ontological analyses were performed on a rank-ordered list of all 288 proteins using Gene Set Enrichment Analysis (GSEA) on GO Biological Processes. Individual nodes are colored by their unadjusted nominal p-value, ranging from highly significant (red) to the significance threshold of 0.05 (blue). The x-axis on the color bar uses a logarithmic scale to capture the range of significance, while bubble size corresponds to the number of proteins (count) associated with each term. Background colored tiles indicate clusters of semantically similar terms. Directional arrows indicate whether terms were up-or-down-regulated. Full results are available in the supplementary materials.

In contrast, KASV infection produced a broader and more sustained proteomic response, predominantly characterized by increased protein abundance at early (3 to 6 DPI) and mid (9 DPI) time points (Fig. 4A), with larger effect sizes and higher statistical significance than observed in MARV infection. Proteins involved in complement activation, coagulation, and endothelial or immune signaling were recurrently among the most significantly affected; these included C4A, MBL2, MASP2, and ICAM1. Liver-derived metabolic enzymes, including ALDOB, were also significantly elevated, consistent with earlier observations of metabolic protein enrichment. Ontological analysis at 6 DPI in KASV infection showed broadly similar downregulation of higher-order cellular processes to that observed in MARV infection, whereas at 9 DPI, proteins associated with signaling and proliferation pathways, including ERK1/2, were enriched among upregulated proteins (Fig. 4C; Supplementary Fig. S7). Direct comparisons between MARV- and KASV-infected bats revealed comparatively few statistically significant differentially abundant proteins, with much of the apparent divergence driven by virus-dependent protein detection rather than shared-protein abundance differences.

Proteins identified as significantly differentially abundant (q ≤ 0.05) in at least one comparison were carried forward for integrated visualization to assess coordinated abundance patterns across all samples. Longitudinal-based clustering of these significant proteins indicated signatures of infection-associated responses (Fig. 5). A prominent cluster showed early and sustained increases in abundance in both MARV- and KASV-infected animals relative to uninfected controls. This cluster was enriched for proteins involved in innate immune activation and complement pathways, including CLU, C4A, C2, MBL2, MASP2, COLEC11, LGALS3BP, ICAM1, and CTSS, many of which ranked among the most significant and highest-magnitude changes in the pairwise DAA. In contrast, a second cluster was comprised of proteins that were consistently decreased during infection, including SERPINA1, SERPINA6, PI16, and CAMP1, reflecting downregulation across multiple timepoints and both viral infections. Related acute phase response proteins (e.g., ALB, AFAM, HP) revealed dynamic longitudinal abundance patterns across both MARV and KASV infections (Supplementary Fig. S6), however, these longitudinal patterns were no longer statistically significant in a multivariate model accounting for sex and technical batch effects.

**Figure 5.**
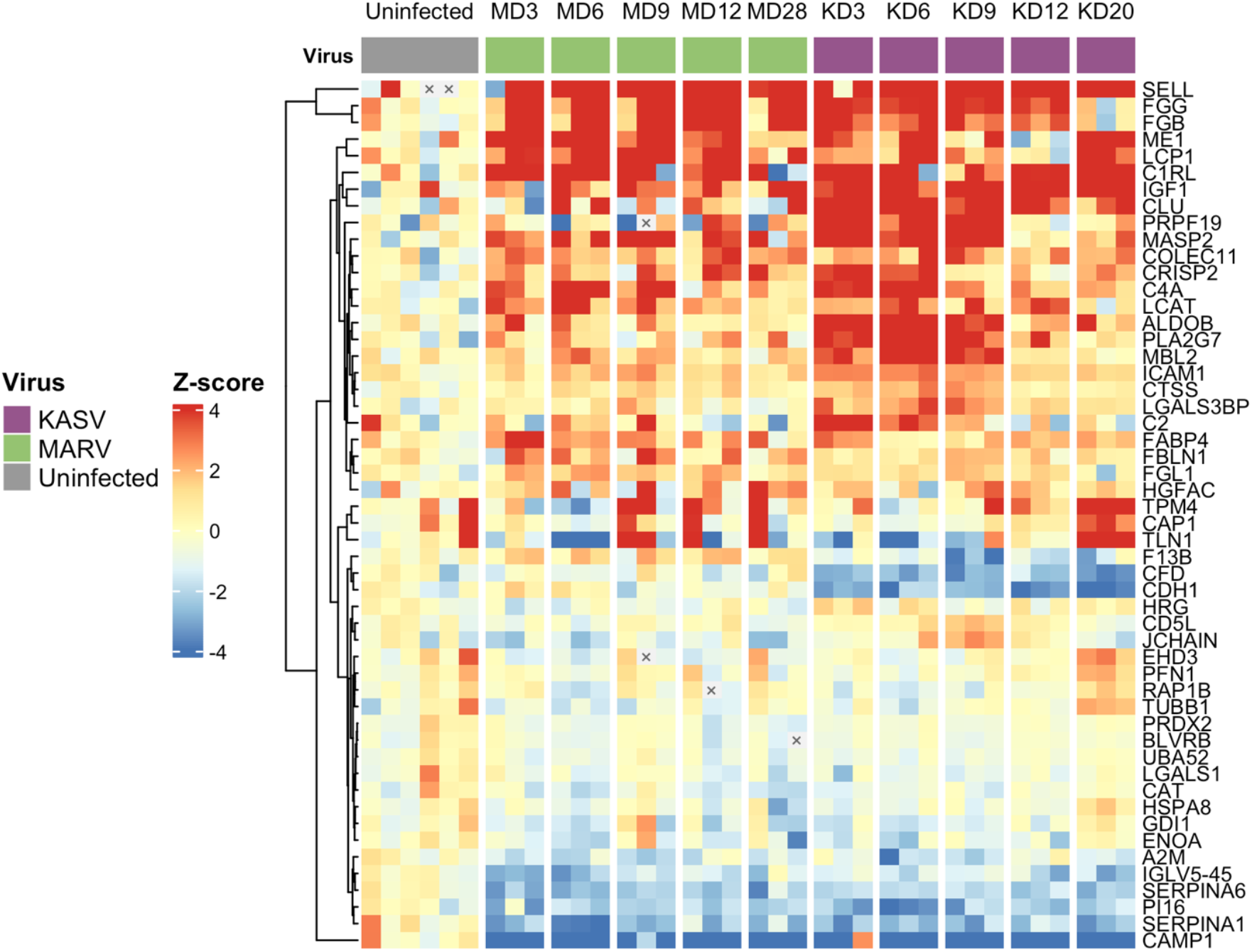
Heatmap of infection-driven changes of the circulating proteome. Heatmap showing z-score–transformed abundances of serum proteins identified as significantly differentially abundant (q < 0.05) in at least one contrast from DAA/volcano plot analyses. Protein intensities were median-centered and scaled relative to uninfected controls using the median absolute deviation, such that values represent the number of standard deviations from the control median. Columns correspond to samples from individual bats ordered by infection group and timepoint (Uninfected, MARV: MD3–MD28; KASV: KD3–KD20; where “D” is shorthand for “days post-infection”), with virus status indicated by the top annotation bar. Rows represent proteins clustered by hierarchical clustering to highlight coordinated abundance patterns. Red indicates increased abundance and blue indicates decreased abundance relative to uninfected controls; grey cells with a black ‘X’ denote missing values.

### Network-based co-abundance analysis reveals virus-dependent module dynamics

To assess coordinated protein behavior beyond individual differential abundance, we performed a network-based analysis of protein co-abundance. Pairwise correlations of protein abundance were used to construct a weighted network, clustered using the Leiden algorithm independently of infection status, time point, or functional annotation. This yielded eight modules of 38 to 68 proteins each (Table 1). Hub proteins, defined by intramodular connectivity (kWithin; mean 0.74 to 0.89), are reported in Table 1. Functional annotation revealed coherent enrichment themes, including protease inhibition, complement activation, proteasome-mediated protein degradation, cytoskeletal organization, and metabolic processes although annotation coverage and statistical significance varied across modules (Supplementary Table 4), with Modules 3–6 showing the most statistically robust annotation across databases.

**Table 1.** Modules of co-varying proteins identified by Leiden-based clustering.

| **Module** | **# Proteins** | **Top Hub Proteins** | **Broad Functional Themes<sup>1</sup>** |
| --- | --- | --- | --- |
| 1 | 64 | SERPINA1 (P01009);<br>SERPINA6 (P08185);<br>ACE (P12821) | Serine protease inhibitor (KW-0722);<br>Signal (KW-0732) |
| 2 | 49 | COLEC11 (Q9BWP8);<br>MASP1 (P48740); LCAT<br>(P04180) | Complement and coagulation cascades<br>(hsa04610); Serpins and Protease<br>inhibitors |
| 3 | 47 | TUBA1B (P68363); TLN1<br>(Q9Y490); MYH9<br>(P35579) | **Autophagy** (R-HSA-9612973);<br>**Phagosome** (hsa04145); **Cytoskeleton**<br>(KW-0206) |
| 4 | 51 | PSMA1 (P25786); PSMA6<br>(P60900); PSMA2<br>(P25787) | **Proteasome** (KW-0647; hsa03050);<br>**Prion and Huntington diseases**<br>(hsa05020; hsa05016); **Negative**<br>**regulation of NOTCH4 signaling**<br>(R-HSA-9604323) |
| 5 | 68 | PLA2G7 (Q13093); CLU<br>(P10909); MBL2 (P11226) | **Metabolic pathways** (hsa01100) |
| 6 | 50 | C1S (P09871); CFHR2<br>(P36980); C1R (P00736) | **Complement and coagulation**<br>**cascades** (hsa04610); **Coronavirus**<br>**disease - COVID-19** (hsa05171);<br>**Signaling by RAS mutants** (R-HSA-6802949); **Common Pathway of Fibrin**<br>**Clot Formation** (R-HSA-140875) |
| 7 | 38 | THBS1 (P07996); CAT<br>(P04040); LDHA (P00338) | Systemic response to oxidative stress;<br>Antioxidant (KW-0049); Oxidoreductase<br>(KW-0560); Neutrophil degranulation (R-HSA-6798695) |
| 8 | 48 | A1BG (P0421); MMP9<br>(P14780); PGLYRP2<br>(Q96PD5) | Regulation of Insulin-like Growth Factor<br>(IGF) transport and uptake by Insulin-like<br>Growth Factor Binding Proteins<br>(IGFBPs) (R-HSA-381426); Extracellular<br>matrix repair |

<sup>1</sup> Modules with statistically significant enrichment terms relative to the study-specific background are indicated in bold. For modules lacking significant study-wide enrichment, results are reported based on a general proteome background to provide functional context; these values are shown in standard font.

To map the global connectivity of the serum proteome, we computed the adjacency between co-abundance modules. Inter-module adjacency, computed by correlating module eigenproteins across all timepoints per virus, revealed consistent positive and negative module–module relationships (Supplementary Fig. S8). Module 4 (Proteasome) and Module 7 (Oxidative Stress/Metabolism) maintained strong positive adjacency in both MARV (r = 0.74) and KASV (r = 0.91), while KASV was associated with stronger negative inter-module relationships overall (e.g., Module 5–Module 3: r = −0.77). Module eigenproteins were further correlated with tissue viral loads stratified by DPI; of 240 tests, few survived FDR correction, though Modules 3, 4, 7, and 8 showed nominal associations at specific infection × tissue × DPI combinations (Fig. 6A). Associations were more pronounced in liver and spleen than blood and were not uniformly shared between viruses.

**Figure 6.**
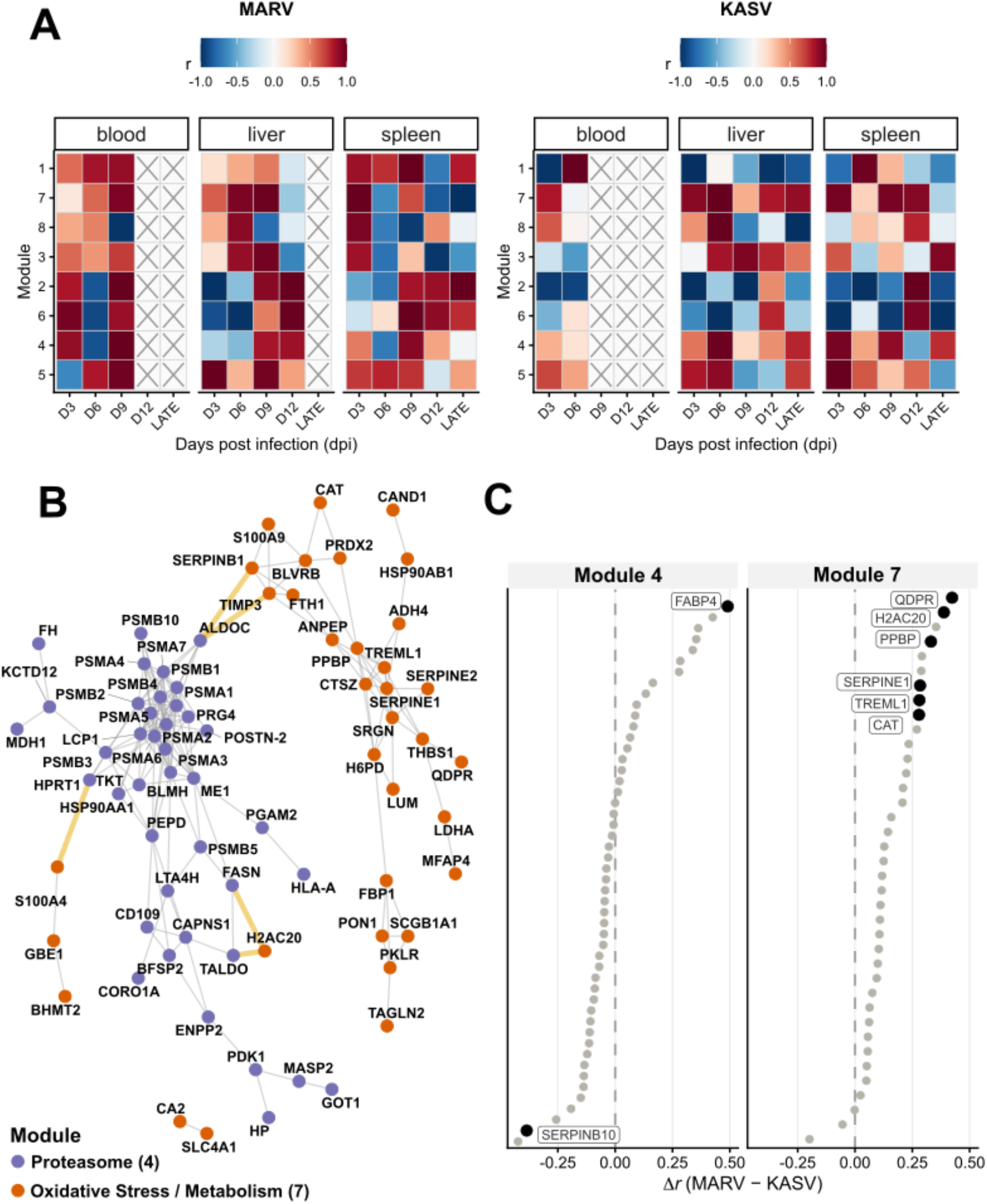
Virus-dependent module dynamics. **A) Module–viral load correlations across tissues and time during MARV and KASV infection**. Heatmaps show Pearson correlations between module-level mean protein expression (z-scored per protein and averaged within modules) and tissue viral loads across days post infection (dpi). Correlations were calculated separately for each virus (MARV, left; KASV, right), time point, and three tissues (blood, liver, spleen). Tile color indicates the correlation coefficient (*r*), with red denoting positive and blue denoting negative correlations. Grey “X” marks indicate module–time–tissue combinations with insufficient data to compute correlations. Modules are ordered by their peak temporal expression, facilitating comparison of virus-specific module–viral load relationships. **B) Inter-Module Network between Module 4 (proteasome-related) and Module 7 (oxidative stress/metabolism).** A focused subnetwork displays the high-confidence (top 200 strongest connections) protein co-abundance correlations between Module 4 (Proteasome; purple) and Module 7 (Metabolic; orange). The layout uses a Fruchterman-Reingold force-directed algorithm to position proteins based on their connectivity, revealing a clear spatial separation of functional clusters that remain linked by a central regulatory core. Inter-module "bridge" connections (highlighted as thick gold lines) are defined as strong correlations ( r > 0.80) between proteins assigned to different Leiden modules. **C) Differential intramodular connectivity of Module 4 and Module 7 proteins between MARV and KASV infection**. Each point represents a single protein, ranked by its differential connectivity score (Δr = r_MARV − r_KASV), where r reflects the mean pairwise Pearson correlation between that protein and all other members of its module, computed separately for each virus cohort. Positive Δr indicates tighter co-abundance with module peers during MARV infection; negative Δr indicates tighter co-abundance during KASV infection. Statistical significance was assessed by permutation testing (n = 1,000 permutations), with FDR correction applied within each module. Black points indicate proteins with nominal significance (permutation p < 0.05); no proteins survived FDR correction, consistent with the limited statistical power of n = 15 animals per virus group.

To examine functional relationships underlying Module 4, we constructed a focused inter-module network of the top 200 co-abundance correlations between Module 4 (Proteasome) and Module 7 (Oxidative Stress/Metabolism; Fig. 6B). Force-directed layout revealed clear spatial separation of proteasome and metabolic clusters connected by eight bridge proteins spanning glycolysis (ALDOC, TALDO, FASN), extracellular matrix remodeling (S100A4, TIMP3, SERPINB1), nucleotide metabolism (HPRT1), and chromatin architecture (H2AC20), with inter-module correlations generally exceeding r > 0.80. To assess whether this coupling was virus-dependent, we quantified differential intramodular connectivity for all Module 4 and 7 proteins using a permutation approach (Fig. 6C). The histone variant H2AC20 — uniquely identified as both a structural bridge protein and the most strongly rewired module member — showed markedly greater co-abundance with module peers during MARV than KASV (Δr = 0.39, p = 0.042), suggesting virus-dependent differences in proteasome–metabolic coupling are concentrated at the inter-module interface. H2AC20 may reflect chromatin-associated protein release or NET-like biology, but this interpretation remains speculative and requires targeted validation.

### Proteasome- and complement-enriched modules exhibit distinct temporal and viral load associations

Because Module 4 was significantly enriched for proteasome-related pathways (KEGG; p-adj = 8.1 x 10-4) as well as pathways related to negative regulation of NOTCH4 signaling (REACTOME; p-adj = 1.8 x 10-3; Table 1) it was examined in greater detail. Eigenprotein trajectories were dynamic during MARV infection through 9 DPI followed by decline, whereas KASV showed a steep drop at 12 DPI with recovery by late-stage infection (Supplementary Fig. S9). Strong multi-module adjacency and a densely connected proteasomal core (PSMB4/PSMA4) linked to metabolic regulators (TKT, ME1; Fig. 6B) position Module 4 as a central, virus-differentially regulated hub.

Module 6 (Complement Cascade) showed a contrasting pattern: during MARV infection it peaked at 6 DPI alongside pronounced negative associations with viral load across all three tissues (Fig. 6A), whereas during KASV infection trajectories peaked later at 12 DPI with strong positive correlations to liver and spleen viral load (Fig. 6A; Supplementary Fig. S9). Complement proteins were distributed across modules in a functionally coherent manner: Module 6 was enriched for pathway initiation and recognition components (C1q, MASP1, C1R, C1S), while Module 3 contained exclusively terminal pathway and membrane attack complex proteins (C5–C7, C8A, C8B, C8G, C9) (Supplementary Table 4).

## Discussion

### Egyptian rousette bats maintain antiviral defense without systemic inflammation

Viral pathogenicity reflects both virus traits and host responses shaped by the evolutionary history of their interaction. In bats, infections with viruses that cause severe disease in humans and NHPs typically produce clinically inapparent outcomes, a pattern often attributed to antiviral resistance or immune tolerance(16, 18, 27). However, growing evidence suggests that these outcomes reflect tightly regulated host responses that limit viral replication while preventing immunopathology rather than absence of inflammation. Our serum proteomic analyses extend previous histological observations by demonstrating that ERBs mount measurable antiviral responses to both MARV and KASV infections while maintaining limited systemic disruption. Changes in circulating proteins were generally modest and transient, peaking during early-to-mid infection (3 to 9 DPI), extending previous transcriptomic findings to the protein level (15, 20). This contrasts sharply with the dysregulated inflammation driving pathology in susceptible hosts, where excessive cytokine signaling (IL-6, IL-1B, TNF, NF-κB/JAK-STAT) causes tissue damage(28–31).

Several features of the ERB serum proteome reinforce this controlled architecture. Apolipoprotein levels remained stable, indicating preserved lipid homeostasis. Complement and innate immune components were detectable at baseline — at qualitatively higher representation than observed in humans(32, 26) — yet infection did not induce large-magnitude acute-phase responses, consistent with the genomic absence of short pentraxins including C-reactive protein (CRP) in ERBs(16). Together, these findings suggest that ERBs maintain a poised antiviral state in which complement and innate defenses are present and functional, but widespread inflammatory escalation is avoided. MARV and KASV differ in their ecological associations with ERBs (a persistent reservoir relationship versus an enzootic cycle involving argasid ticks) as well as in replication kinetics and tissue tropism. These differences provide a plausible biological basis for the virus-dependent serum proteomic signatures we observed, though ecological or evolutionary history alone is unlikely to fully explain them.

### Virus-dependent proteomic signatures reflect divergent infection dynamics and distinct liver pathophysiology

Although MARV and KASV elicited broadly overlapping responses, their magnitude, timing, and composition differed substantially. MARV induced transient perturbations peaking at 6 DPI, including the upregulation of acute-phase proteins (SERPINA6, SERPINA1, COLEC11, C4A, ICAM1) and downregulation of oxidoreductases (BLVRB, PRDX1, CAT) that resolved by mid-infection, consistent with efficient viral control. KASV produced a broader, more sustained signature including repeated detection of complement components (C4A, C2, CLU), lectin pathway proteins (MBL2, MASP2), and multiple liver-associated metabolic enzymes (ALDOB, BHMT, AKR1A1, ALDH1A1, SORD) detected exclusively in KASV-infected animals. These findings align with prior reports demonstrating stronger early hepatic replication and transient lymphohistiocytic hepatitis during KASV infection(8, 33, 13). Sorbitol dehydrogenase (SORD), a highly specific marker of hepatocellular injury in humans(34), peaked at 9 DPI, suggesting subclinical membrane disruption not always captured by conventional clinical chemistry panels. These findings indicate that KASV engages complement and alters hepatic homeostasis more extensively than MARV while still resolving without overt systemic disease. The detection of these subtle, virus-dependent signatures (several of which likely reflect post-translational or protein-level regulation) underscores the potential of proteomic approaches to reveal infection dynamics that may be less apparent at the transcript-level alone.

Network analyses reinforced these distinctions. Complement-associated Module 6 peaked at 6 DPI and negatively correlated with MARV viral load, whereas during KASV infection it peaked later (12 DPI) with positive correlations to liver and spleen viral burden. This module is functionally enriched for pathways associated with hyperinflammatory responses in humans, particularly those observed during SARS-CoV-2 infection. These opposing dynamics suggest that shared functional modules are deployed under distinct regulatory constraints depending on the infecting virus, emphasizing the utility of these proteomic approaches to evaluate systems-level coordination beyond single-protein effects.

### Proteasome-associated antiviral responses

A prominent systems-level feature was coordinated regulation of proteasome-associated proteins (Module 4). Multiple lines of evidence support regulated remodeling rather than nonspecific protein turnover: immunoproteasome subunits were detected in mature, proteolytically processed forms, supporting catalytically competent complexes, while concurrent detection of constitutive PSMB7 across time points and individual bats suggests preserved 20S barrel integrity alongside inducible subunit enrichment(35). Module 4 dynamics were temporally structured and virus-dependent, mirroring differences observed in complement modules and their relationships to viral load.

Network topology revealed tight coupling between Module 4 (Proteasome) and Module 7 (Oxidative Stress/Metabolism) via eight bridge proteins spanning glycolysis, lipid metabolism, extracellular remodeling, and inflammatory regulation. For example, Fatty Acid Synthase (FASN) and Tissue Inhibitor of Metalloproteinases 3 (TIMP3) exemplify functional intersections between metabolism, proteostasis, and host–virus interactions observed in other mammals. FASN, a de novo fatty acid synthase, may reflect hepatocellular metabolic activity supporting energy production and protein turnover, processes that can also intersect with viral replication(36). TIMP3, an extracellular matrix regulator, may reflect mechanisms of inflammatory restraint: in other mammals, cell-surface TIMP3 is dynamically controlled via LRP1-mediated endocytosis and shedding, modulating the timing of TNF release(37). While these mechanistic links are speculative, they provide a conceptual framework for how ERBs might limit tissue-damaging inflammation while maintaining effective antiviral responses.

H2AC20, a bridge protein with strong differential connectivity, is a core histone variant whose extracellular detection in serum is increasingly recognized as a marker of neutrophil extracellular trap (NET) formation in humans(38). NETs are chromatin-based antimicrobial structures decorated with histones, myeloperoxidase, and neutrophil elastase, implicated in antiviral defense across a range of pathogens. Elevated circulating H2AC20 and other NET-associated markers have been linked to early cardiac damage in severe COVID-19(39), reflecting the intensity of innate inflammatory responses. The stronger MARV-associated co-abundance of H2AC20 with metabolic and proteasomal module peers may therefore reflect a coordinated, regulated NET-like response more tightly integrated into the systemic proteomic signature during MARV than KASV infection. Whether this represents genuine NETosis, passive histone release, or a bat-specific variant of this process remains to be determined but raises the intriguing possibility that ERBs deploy chromatin-based innate mechanisms in a virus-dependent manner without the immunopathological consequences observed in humans.

In mammalian systems, acute infection drives IFN-γ–dependent proteasome remodeling, incorporating inducible subunits (β1i, β2i, β5i) that enhance MHC class I peptide generation(40–42). IFN-γ signaling appears species-specific in ERBs, robustly inducing interferon-stimulated genes in ERB cells but minimally in comparable human systems(43). Comparative genomics further reveals distinctive MHC-I features in fruit bats, potentially shaping the presented peptide repertoire(16, 44). Elevated baseline immunoproteasome representation may therefore prime hepatocytes for rapid antigen processing upon infection while limiting high-magnitude inflammatory induction(41, 45). During MARV infection, relative Module 4 stability coupled with a mid-infection Module 7 rise may reflect controlled resource mobilization and subsequent resolution. The late-stage Module 4 decline during KASV infection — coinciding with broader hepatic engagement and lacking synchronized metabolic recovery — aligns with histopathological evidence that KASV induces more pronounced hepatic involvement than MARV(13). We propose these patterns may reflect a shift from coordinated antigen processing during MARV to prolonged hepatic stress during KASV. Importantly, these interpretations remain theoretical and require targeted experimental validation. Future work could quantify immunoproteasome assembly in ERB hepatocytes – for example, by purifying 20S complexes after IFN-γ stimulation – to assess functional incorporation of inducible subunits.

### Multilayered analysis reveals biology not apparent from differential abundance alone

A key strength of this study is its multilayered analytical framework. Network-based analysis identified coordinated protein communities, including regulatory hubs and bridge proteins structuring immune and metabolic pathways, that were not apparent from single-protein statistics and were not among the most differentially abundant proteins by conventional thresholds. Interpretability was strengthened by the use of captive-reared, experimentally infected ERBs with documented life histories, reducing background ecological and demographic variability. Both infecting viruses were low passage bat and tick-derived isolates (371bat MARV; UGA-Tick-20170128), reducing potential confounding effects from cell culture adaptation. Integration of serum proteomic data with previously published tissue-level viral replication data from the same animals(8, 10) enabled biologically grounded interpretation of systemic signatures. Convergence across differential abundance, network structure, and prior virological data increases confidence that observed patterns reflect coordinated host programs rather than stochastic variation.

### Limitations

Serum proteomics provides a snapshot of the circulating proteome but does not directly resolve tissue of origin. Although integration with prior tissue viral load data strengthens inference regarding hepatic involvement, direct quantification of proteasome regulation and complement activation within specific organs will be necessary to confirm the proposed mechanisms. In addition, network inference identifies coordinated patterns but does not establish causality(46, 47). Module associations with viral load and infection stage should therefore be interpreted as correlative rather than mechanistic.

Study design constraints also warrant consideration. Sample sizes were limited, and the cross-sectional design (using age-matched uninfected bats rather than paired within-individual longitudinal sampling) constrains resolution of intra-individual trajectories, though it enabled use of well-characterized archival specimens while minimizing animal use. Finally, captive experimental infections reduce environmental influences but may not fully reflect the ecological complexity of natural exposures, including repeated infection, coinfection, nutritional variability, and stress-related immune modulation(14, 48–50). Despite these limitations, the convergence of differential abundance, network structure, and prior virological data supports the conclusion that ERBs deploy tightly regulated, virus-dependent systemic responses that balance complement activation and metabolic control without progressing to overt pathology.

## Materials and Methods

### Bats and Biosafety

Experiments with animals and viable Marburg virus (MARV) complied with all relevant regulations, with study protocols (#2977, #3090) overseen and approved by CDC’s Institutional Animal Care & Use Committee (IACUC), Animal Care and Use Program Office (ACUPO), Comparative Medicine Branch (CMB), and Institutional Biosecurity Board (IBB), using guidelines established by the Association for the Assessment and Accreditation of Laboratory Animal Care, International (AAALAC), the Animal Welfare Act and Regulations, and The Guide for the Care & Use of Laboratory Animals(8, 10, 15, 53). All investigators and animal handlers followed strict BSL-4 biosafety and infection control practices documented in Amman et al. 2015, Schuh et al. 2017, and Schuh et al. 2022. Fresh fruit was provided daily, and water was provided ad libitum.

The uninfected cohort consisted of male (n=3) and female (n=3) originating from an established, MARV-naïve, multi-generational, captive colony(26).

### Ethics Statement

Work with ERBs was approved by the Institutional Animal Care and Use Committee (IACUC) of the CDC. The CDC is accredited by the Association for Assessment and Accreditation of Laboratory Animal Care International (AAALAC).

### Experimental Design, Specimen Collection and Sample Testing

Detailed study protocols for each infection cohort (specimen collection schedule and techniques), collection of virological and clinicopathologic samples and data, and the initial histopathological characterizations from these studies were previously described, in detail, in Amman et al. 2015 (MARV) and Schuh et al. 2022 (KASV). Brief descriptions of the study methods are as follows:

MARV: All bats were anesthetized with isoflurane and inoculated subcutaneously with injection at the mid-ventral abdomen at 0 DPI. Bats received 4 x log10TCID50 of the MARV-371bat virus (second passage on Vero-E6 cells; Genbank: FJ750958;Towner et al. 2009) strain of MARV diluted in a solution of Dulbecco’s modified Eagle’s medium (DMEM). Blood samples were collected via venipuncture from the cephalic vein. All blood and tissues collected at necropsy were analyzed with quantitative reverse transcriptase PCR (qRT-PCR) targeting VP40 and using reagents and procedures described by Amman et al. (2012)(11). Aliquots of whole blood were used to monitor for viremia, and antibody responses were assessed by enzyme-linked immunosorbent assay (ELISA) for IgG antibodies reactive to MARV. All bats were serially euthanized (three bats per time point obtained from the initial infection study(10)) by an overdose of isoflurane followed by cardiac exsanguination.

KASV: To mimic a tick bite, all bats were inoculated by the intradermal route in the subcaudal abdominal region under isoflurane anesthesia at 0 DPI. Bats received 4 x log10TCID50 of the UGA-Tick-20170128 strain of KASV (GenBank: MT309090, MT309094, and MT309097) prepared in 0.1 mL of sterile PBS. Blood samples were collected via venipuncture from the cephalic vein using a sterile lancet. Aliquots of whole blood were used to monitor for viremia (KASV RNA by qRT-PCR) and antibody responses (anti-KASV IgG by indirect ELISA). Tissues collected at necropsy from all bats were tested for KASV RNA by qRT-PCR to determine virus-tissue tropism. All bats were serially euthanized (three bats per time point obtained from the initial infection study(8)) by an overdose of isoflurane followed by cardiac exsanguination.

Uninfected: Study and sample collection protocols for the uninfected bat group are detailed in Genovese et al. 2026(26). Briefly, uninfected bats were age- and sex-matched adults, and all female bats were reproductively mature but confirmed not pregnant at the time of specimen collection. Blood samples were collected by venipuncture of the cephalic vein on the propatagium using a sterile blood lancet (Premiere #95-7820, C&A Scientific, Manassas, VA, USA). Serum was separated from whole blood by centrifugation at 12000 RCF for 90 seconds before inactivation procedures.

### Virus inactivation procedures

All specimens used in this study (including uninfected specimens) received a dose of 5 megarads of gamma-radiation while on dry ice(54, 55).

### Serum Sample Preparation

For the digestions, all inactivated serum specimens were thawed to room temperature on wet ice, spun down, and then 5 μl of each serum sample was aliquoted into prepped Eppendorf tubes in 5 % sodium dodecyl sulfate (SDS), 50 mmol/L triethylammonium bicarbonate (TEAB) buffer, and LCMS-grade water. Proteins were digested via suspension-trap devices (S-Trap; ProtiFi, 100 to 300 μg binding capacity). The S-Trap is a powerful Filter-Aided sample preparation (FASP) method that traps acid aggregated proteins in a quartz filter prior to enzymatic proteolysis and allows for reduction/alkylation/tryptic proteolysis all in one vessel. The enzymatic digestion was initiated with a first addition of trypsin 1:100 enzyme:protein (wt/wt) for 4 hours at 37°C, followed by a boost addition of trypsin using same wt/wt ratios for overnight digestion at 37 °C. Peptides were eluted from the S-Trap with sequential elution buffers of 100 mmol/L TEAB, 0.5 % formic acid, and 50 % acetonitrile (ACN) 0.1 % formic acid (percents are volume ratios). The eluted tryptic peptides were dried in a vacuum centrifuge and re-constituted in 0.1 % trifluoroacetic acid. These were subjected to liquid chromatography mass spectrometry (LC-MS) analysis.

### Liquid Chromatography-Mass Spectrometry (LC-MS) Analysis of Serum Samples

Peptides were resolved on a Thermo Scientific Dionex UltiMate 3000 RSLC system: PepSep 150 µm x 25 cm C18 column (PepSep; Denmark) with 1.5 μm particle size (100 Å pores), heated to 40 °C. A volume of 5 μl was injected corresponding to 1 μg of total peptide, and separation was performed in a total run time of 90 min with mobile phases A: 0.1 % formic acid in water and B: 80 %ACN, 0.1 % formic acid. Separated peptides were electrosprayed directly into a tribrid Fusion Lumos mass spectrometer (Thermo Fisher Scientific), operated in data-independent acquisition mode (DIA). A survey full scan MS (from m/z 350 to 1200) was acquired in the Orbitrap at a resolution of 120,000 (at 200 m/z). For the fragmentation spectra, the following settings were used: the mass range of 350 to 1200 Da was segmented into 19 windows, overlapping by 1 Da, with an isolation window width of 45.7 Da. They were fragmented via higher collisionally-induced dissociation (HCD) at 33 % normalized collision energy. Automatic gain control (AGC) target set to 1000 % (5 x 105) and ion filling time was set to automatic. Detection was in the Orbitrap at a resolution of 15,000.

### Data Processing with Spectonaut

Mass spectrometry raw data files were processed with Spectronaut version 18 (Biognosys, Zurich, Switzerland) using DirectDIA analysis mode, with a decoy FDR at less than 1% for peptide spectrum matches and protein group identifications was used for spectra filtering (Spectronaut default). For downstream analysis, a report was generated and exported from Spectronaut using the scheme provided by Mass Spectrometry Downstream Analysis Pipeline (MS-DAP; version 1.0.6)(56). The quantitative report exported from Spectronaut was processed using the MS-DAP R package, which was also used for preprocessing and quality control of this dataset prior to differential expression/abundance and network analyses. Detailed search parameters are provided in the supplemental methods.

### Protein Ortholog Mapping

Bat protein identifiers from Spectronaut/MS-DAP DIA analysis were mapped to human orthologs via BLASTp (NCBI BLAST+ v2.14.0, implemented through rBLAST v0.99.3) against the reviewed human Swiss-Prot proteome (UniProt, 09-2025). Top hits per query were selected using a custom sorting strategy as described in Genovese et al. 2026(26), with manual verification/correction of unmatched or misannotated entries. Details about database construction, query parameters, and manual annotation records are provided in Supplemental Methods and the supplemental protein list.

### Differential abundance and detection analysis

Differential expression and detection analyses were performed in MS-DAP using the DEqMS algorithm, which accounts for variance dependence on the number of quantified peptides (i.e. PSM counts) per protein, using log-transformed and normalized protein abundance values(57). Proteins were required to have ≥1 peptide with confidence ≥0.01 in ≥75% of samples per group to be tested. Statistical contrasts (CTRL vs. infection × dpi groups) were modeled with linear regression adjusting for sex and batch, with significance defined as q < 0.05. Full contrast definitions and metadata are provided in Supplemental Methods.

### Protein Co-abundance Network Analysis

Protein co-abundance networks were constructed by adapting the transcriptomic workflow “Simple Tidy GeneCoEx” described by Li et al. (2023)(58) for proteomic data. Gene co-expression analysis is an effective approach to detect modules of co-expressed genes (or proteins) that display similar expression patterns and may function in related biological processes. For network analysis, missing values were imputed per protein using half the minimum non-zero abundance observed for that protein (log2 scale), and duplicate protein entries were resolved by averaging abundances and retaining a single merged entry. Full imputation and deduplication procedures are documented in Supplemental Methods.

## Acknowledgments

The authors would like to thank the researchers and staff at the UC Davis One Health Institute, the UC Davis Proteomics Core Facility, the Centers for Disease Control and Prevention – Viral Special Pathogens Branch, and the National Institute of Standards and Technology for their administrative and scientific support for the work. The findings and conclusions in this report are those of the authors and do not necessarily represent the official position of the Centers for Disease Control and Prevention. We would also like to acknowledge and thank Eunah Preston at the UC Davis One Health Institute for her assistance in designing schematic figures and visualizations for this report.

Identification of certain commercial equipment, instruments, software, or materials does not imply recommendation or endorsement by the National Institute of Standards and Technology, or the Centers for Disease Control and Prevention, nor does it imply that the products identified are necessarily the best available for the purpose. These opinions, recommendations, findings, and conclusions do not necessarily reflect the views or policies of NIST or the United States Government.

## Data availability

The mass spectrometry proteomics data have been deposited to the ProteomeXchange Consortium via the PRIDE partner repository with the dataset identifier PXD082683.

## Author Contributions

The manuscript was written through contributions of all authors. All authors have given approval to the final version of the manuscript. Conceptualization: B.N.G., S.J.A., J.K.M., J.S.T., and B.H.B.; Technical Study Design: B.N.G., G.G., J.K.M. and B.H.B.; Data Collection: B.N.G., G.G., A.J.S., B.R.A., J.A.E.; Data Analysis: B.N.G., N.R., G.G.; Data Interpretation: B.N.G., N.R., B.N., S.J.A., J.K.M., J.S.T., B.H.B.; Data Visualization: B.N.G., N.R.; J.K.M., B.H.B.; Resources: J.S.T.; Funding Acquisition: B.N.G., J.K.M., B.H.B.; Writing – original draft: B.N.G.; Writing – review and editing: all authors

