## Supplemental Figures for "Proteomic comparison of Marburg and Kasokero virus infection in natural host Egyptian rousette bat reveals distinct antiviral pathways"

**Supplementary Materials**

**S1: Principal Component Analysis (PCA) plot by day post-infection.**

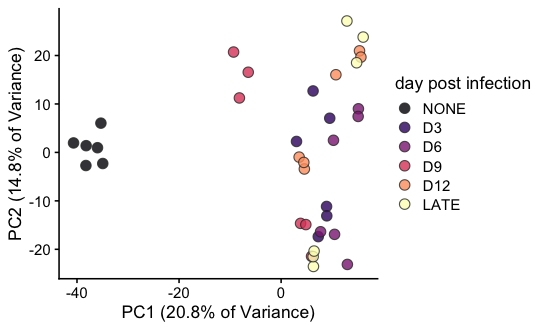

**S1: Principal Component Analysis (PCA) plot by day post-infection.** The PCA plot shows protein profiles colored by day post-infection (DPI). The ‘LATE’ time point represents the latest DPI collected for each experimental cohort (MARV = 28 DPI; KASV = 20 DPI).

**S2: Variance Partition Analysis.**

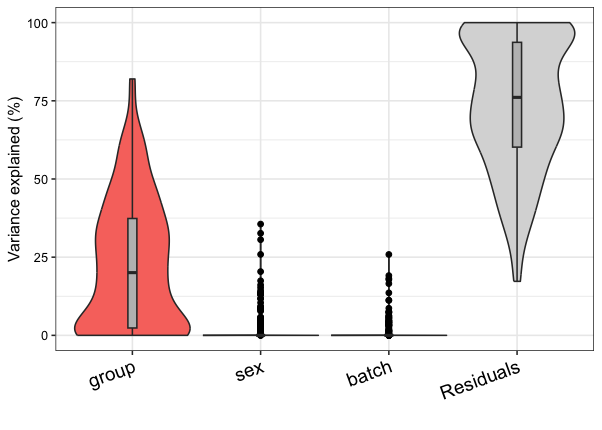

| **Statistic** | **Group (infection-dpi)** | **Sex** | **Batch** | **Residuals** |
| --- | --- | --- | --- | --- |
| **Boxplot Upper Whisker** | 82.0 | 0.0 | 0.0 | 100.0 |
| **Boxplot Upper Hinge** | 37.4 | 0.0 | 0.0 | 93.7 |
| **Mean** | 23.1 | 1.8 | 1.1 | 74.1 |
| **Median** | 20.1 | 0.0 | 0.0 | 76.1 |
| **Boxplot Lower Hinge** | 2.3 | 0.0 | 0.0 | 60.2 |
| **Boxplot Upper Whisker** | 0.0 | 0.0 | 0.0 | 17.3 |

**S2: Variance Partition Analysis.** Variance partitioning analysis was performed to quantify the contribution of different experimental factors to overall protein abundance variation. The analysis evaluated three factors: group (experimental condition), sex, and batch effects. The majority of variance remains unexplained by the model (mean residuals: 74.1 %), which is typical for proteomic datasets and reflects a combination of biological variability and technical noise. The distribution of variance explained by group shows considerable heterogeneity across proteins (interquartile range: 2.3 to 37.4 %), suggesting that while the experimental grouping strongly affects many proteins, others are less responsive to the conditions tested.

**S3: Serum trajectories of 10 metabolic proteins exclusively detected in KASV-infected samples.**

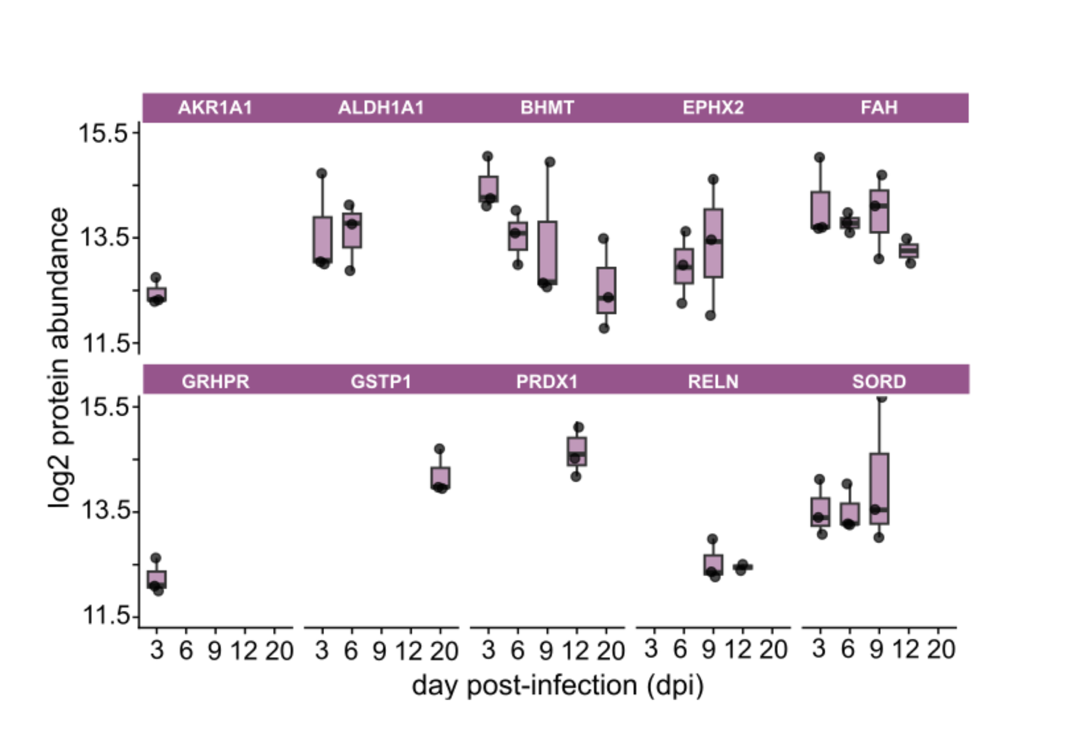

**S3: Serum trajectories of 10 metabolic proteins exclusively detected in KASV-infected samples.** The y-xis represents log2 protein abundance (unimputed) and the x-axis represents day post-infection with KASV. There are three individual bats per timepoint, and only proteins detected in all bats at at least one timepoint are shown. **Abbreviations:** Aldo-keto reductase family 1 member A1 **(AKR1A1);** Aldehyde dehydrogenase 1A1 **(ALDH1A1);** Betaine--homocysteine S-methyltransferase 1 (**BHMT);** Bifunctional epoxide hydrolase 2 **(EPHX2);** Fumarylacetoacetase **(FAH);** Glyoxylate reductase/hydroxypyruvate reductase **(GRHPR);** Glutathione S-transferase P **(GSTP1);** Peroxiredoxin-1 **(PRDX1);** Reelin **(RELN);** Sorbitol dehydrogenase **(SORD).**

**S4: Detection of 40 shared proteins across infection groups (MARV and KASV)**

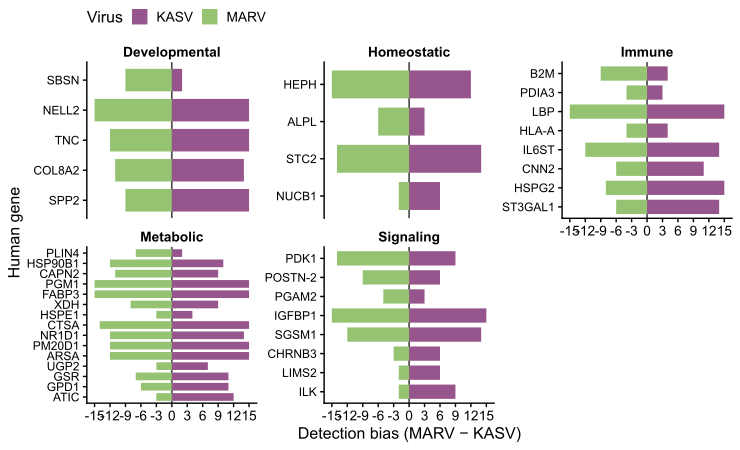

**S4. Detection of 40 shared proteins across infection groups (MARV and KASV).** Diverging bar plots show the number of samples (out of 15 per infection group) in which each protein was detected during MARV (left) or KASV (right) infection. Proteins are ordered within broad biological functional annotation categories by detection (MARV − KASV). The vertical line marks zero bias. Proteins (y-axis) are represented by their orthogonal human gene symbol, and the x-axis indicates protein counts.

**S5: Sequence comparison of two CAMP-like proteins.**

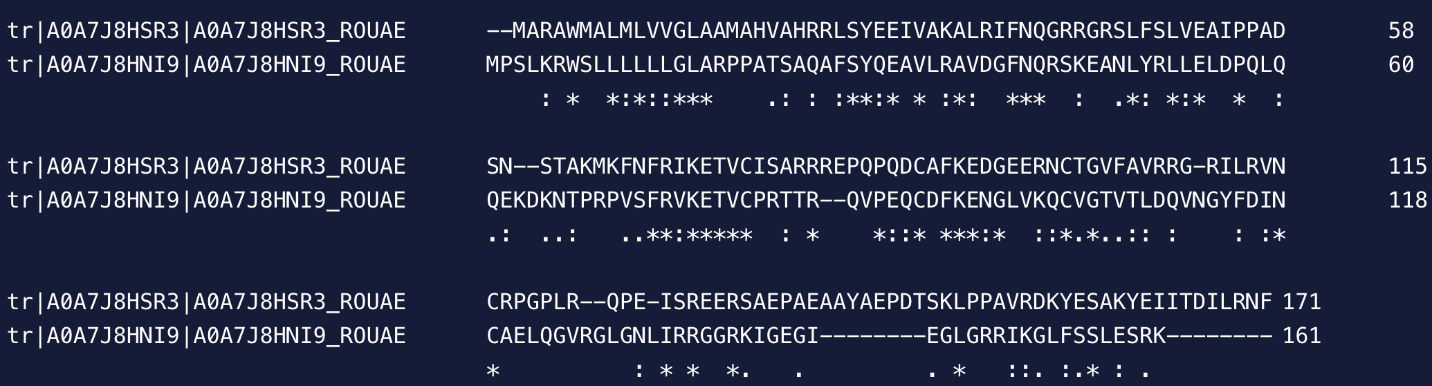

**S5: Sequence comparison of two CAMP-like proteins.** Multiple sequence alignment of two *Rousettus aegyptiacus* proteins (A0A7J8HSR3 and A0A7J8HNI9) showing conserved motifs alongside N-terminal differences, consistent with them being related paralogs rather than alternative splice variants. Highly conserved residues are indicated by asterisks (*), strongly similar residues by colons (:), and weakly similar residues by periods (.), indicating stretches of shared sequence as well as insertions/deletions that may reflect functional divergence. Alignment was conducted using CLUSTAL (1.2.4) via UniProt.

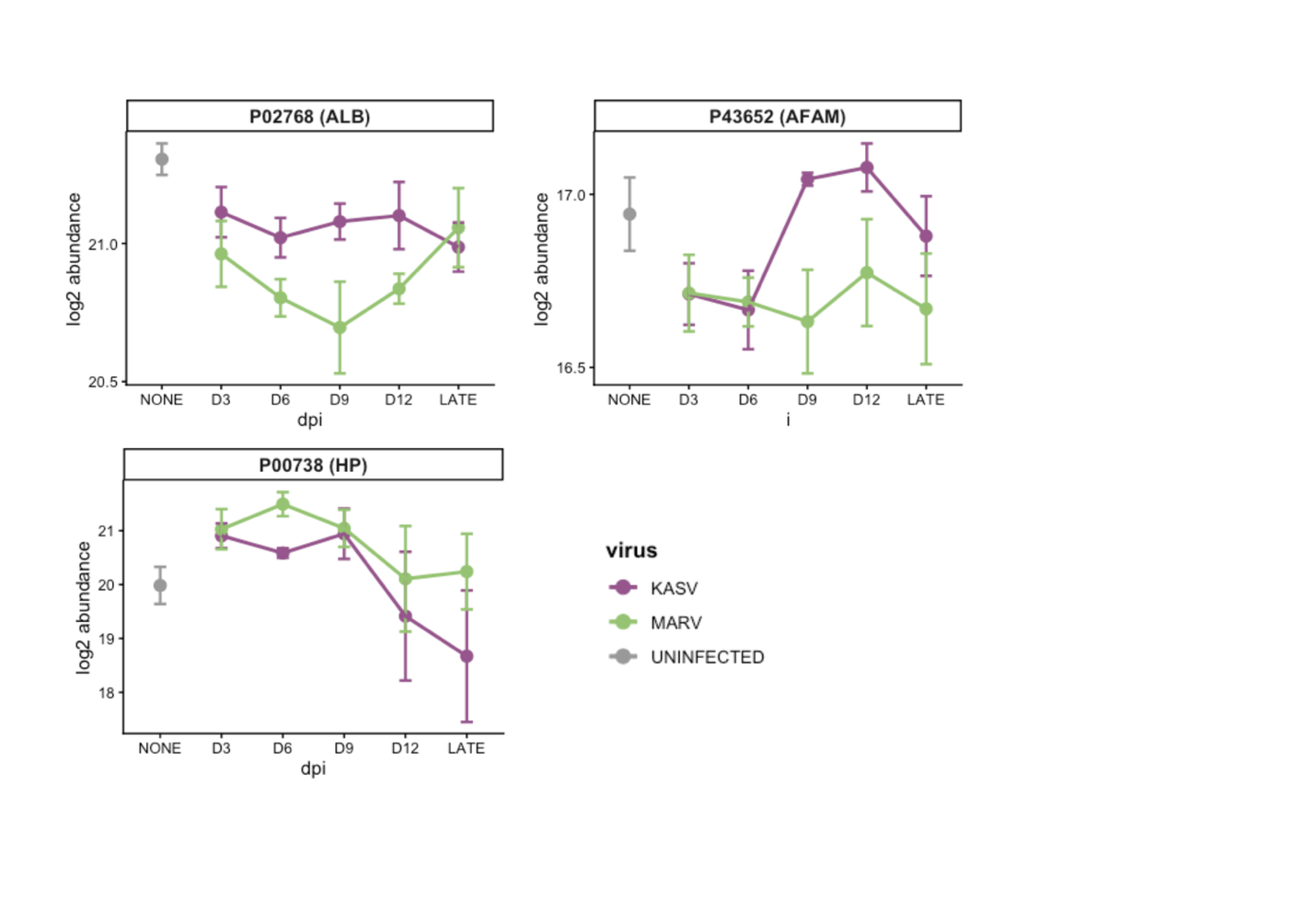
**S6: Abundance plots for Albumin (ALB), Afamin (AFAM), and Haptoglobin (HP).**

**S6. Abundance plots for Albumin (ALB), Afamin (AFAM), and Haptoglobin (HP).** Line plots represent log2 abundance values of five key proteins identified in serum specimens from uninfected (grey), KASV-infected (purple), and MARV-infected (green) ERBs. Data points represent the mean abundance at each day post-infection (DPI), with error bars indicating the standard error of the mean. Bat proteins are listed by the UniProt ID of the corresponding human ortholog alongside the human gene symbol.

**S7: Preliminary GO:BP enrichment analysis of significantly differentially abundant proteins (Uninfected vs. KASV - 9 DPI)**

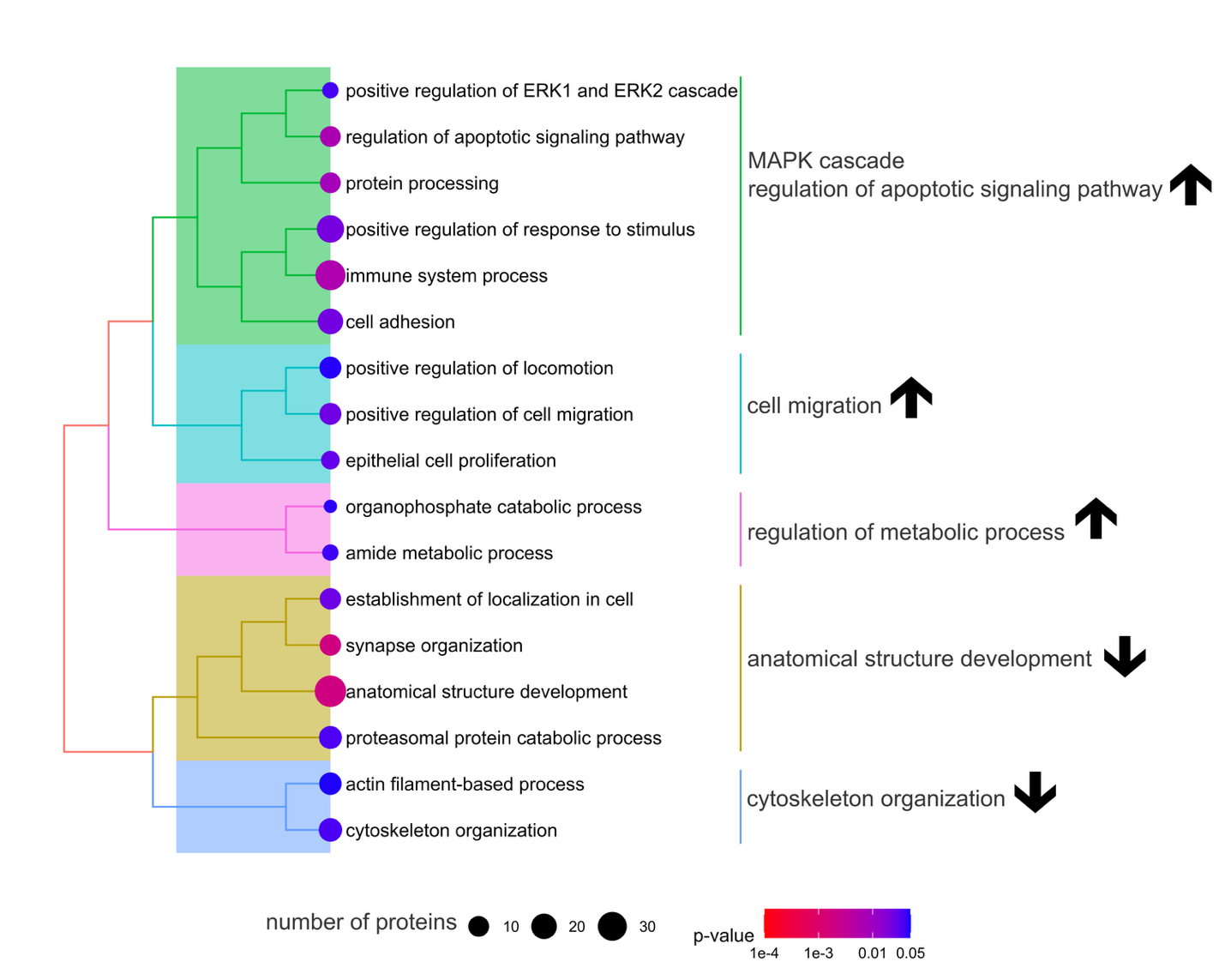

**S7: Preliminary pathway and process enrichment analysis of significantly differentially abundant proteins (Uninfected vs. KASV – 9 DPI).** The treeplot illustrates enriched terms for differentially abundant proteins in Uninfected vs. KASV (9 DPI) comparison.

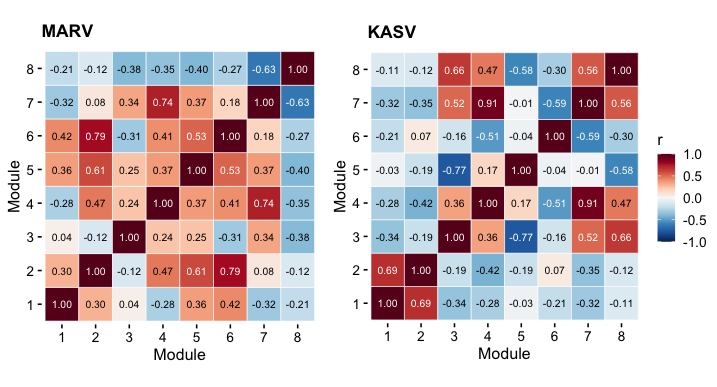
**S8: Comparative inter-module adjacency matrices.**

**S8. Comparative inter-module adjacency matrices.** Heatmaps display the Pearson correlation coefficients (r) between module eigenproteins for MARV (left) and KASV (right) infections. Modules were defined using a stable Leiden clustering algorithm. Positive correlations (red) indicate synchronized abundance patterns between modules, while negative correlations (blue) reflect divergent temporal trajectories of module proteins. All modules (1 to 8) correspond to the same protein clusters identified in the global temporal correlation and trait association analyses.

**S9: Temporal abundance trajectories of selected protein modules during MARV and KASV infection.**

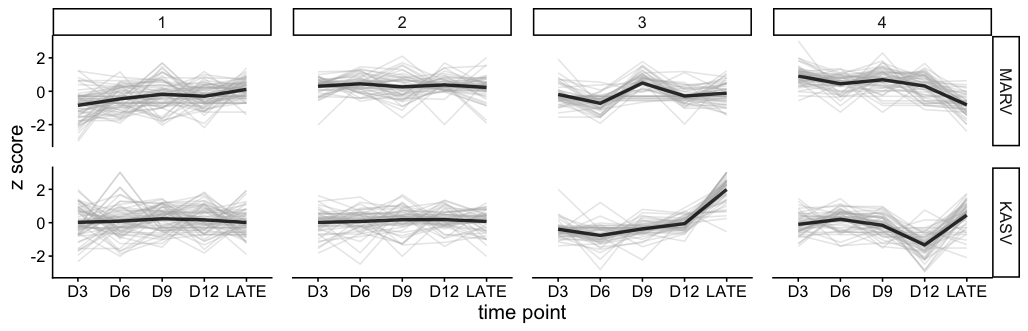

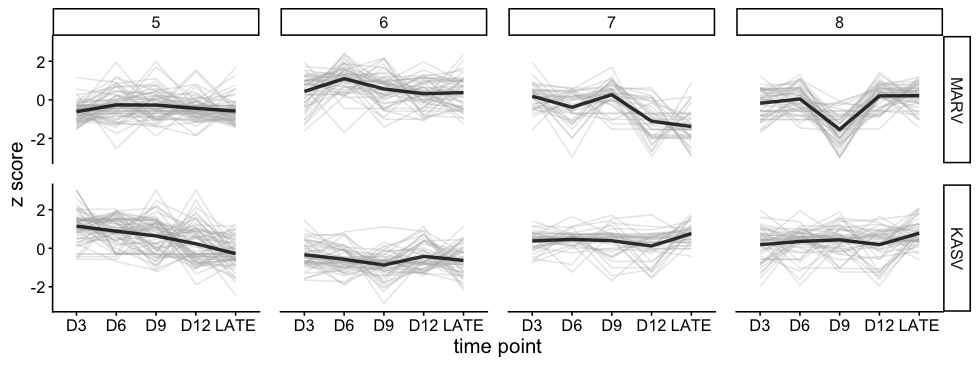

**S9. Temporal abundance trajectories of selected protein modules during MARV and KASV infection**. Line plots show z-scored protein abundances across days post infection (D3, D6, D9, D12, LATE) for five Leiden-derived co-abundance modules, Each panel corresponds to a single module (columns) and virus condition (rows: MARV or KASV). Thin grey lines represent individual proteins within each module, while the thicker colored line indicates the module mean z-score at each time point, summarizing the coordinated temporal behavior of module members. Z-scores were calculated per protein across samples prior to module aggregation. A drop or rise in module z-score reflects a coordinated relative decrease or increase of proteins in that module, compared to their own average behavior across conditions.

**S10: Resolution parameter optimization for Leiden Clustering**

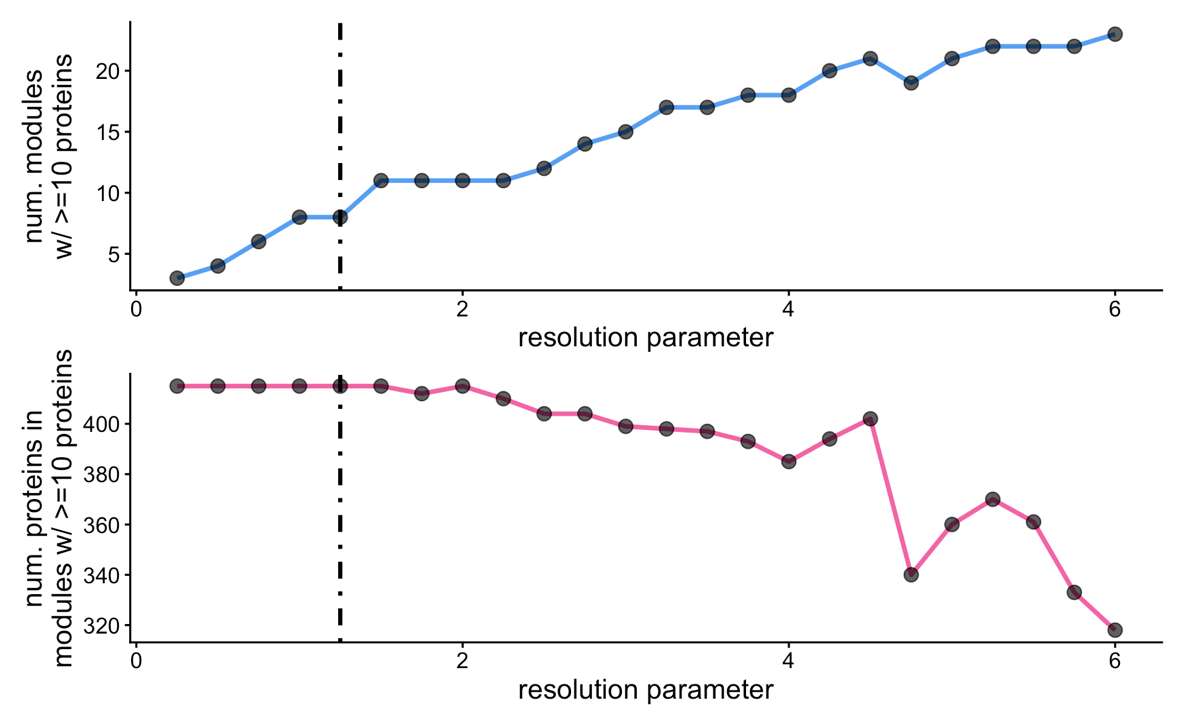

| **Number of Modules** | **Number of Proteins Contained** | **Resolution Parameter** |
| --- | --- | --- |
| 3 | 415 | 0.25 |
| 4 | 415 | 0.50 |
| 6 | 415 | 0.75 |
| 8 | 415 | 1.00 |
| **8** | **415** | **1.25** |
| 11 | 415 | 1.50 |
| 11 | 412 | 1.75 |

**S10. Resolution parameter optimization for Leiden Clustering.** Multi-panel line plots illustrating the impact of varying the resolution parameter (range: 0.25 to 6.0) on network modularity. The top panel tracks the number of robust modules containing at least 10 proteins, showing a stable plateau of 8 modules between resolution 1.0 and 1.25. The bottom panel displays the total number of proteins contained within these robust modules; a resolution of 1.25 was selected (indicated by the dashed vertical line) as it maximized the number of stable modules while maintaining protein inclusion (415 proteins).
